# A state-structured pharmacodynamic framework separating resistance, tolerance and persistence, applied to Mycobacterium tuberculosis and Staphylococcus aureus

**DOI:** 10.1101/2025.02.12.637810

**Authors:** Vahhab Piranfar

## Abstract

Antibiotic resistance, tolerance, and persistence represent key bacterial survival strategies that impact treatment outcomes and global health. While *Staphylococcus aureus* is a rapidly growing pathogen associated with acute infections, *Mycobacterium tuberculosis* exhibits slow growth and chronic persistence, necessitating prolonged antibiotic regimens. In this study, we developed a state-structured pharmacodynamic model in which replicating and dormant subpopulations each carry their own concentration-response, so that the three strategies separate onto distinct measurable axes: resistance shifts the minimum inhibitory concentration, tolerance multiplies the minimum duration for killing, and persistence lifts the deep killing endpoint alone. Using global variance-based sensitivity analysis and profile likelihood, we identified which parameter governs the length of therapy and which parameters can be estimated at all from time-kill data. Our findings show that in the slow-growing organism 87% of the first-order variance in time to sterilisation is carried by the rate at which dormant cells resume replication, rather than by the rate at which dormant cells are killed, placing resuscitation at the centre of regimen shortening and identifying a class of intervention worth measuring. This version supersedes version 1, whose closed-form biphasic killing law is discontinuous and whose quantitative results are withdrawn and corrected here; the work is a modelling and methods contribution, contains no experimental data, and its parameter values are illustrative rather than measured.

**Summary of changes in version 2:** This version corrects errors in version 1 that alter its conclusions. Readers of version 1 should treat its quantitative results as withdrawn.

*Mathematics:* The biphasic killing law of version 1 (its Equation 4) is discontinuous: at the transition time its second branch evaluates to exactly the inoculum, so the modelled population rises by 2.6 to 3.5 log10 at the breakpoint. It also converges to a permanent floor, making sterilisation impossible at any duration. Its transition time is not an independent parameter and, on time-kill data of realistic density, is not estimable at all. The resistance equation omitted the carrying capacity declared alongside it and reached 10^57 CFU/mL by day 10. The tolerance equation omitted replication, which reverses the sign of its result for *S. aureus*: version 1 reported fifteen logs of killing where its own parameters imply net growth.

*Withdrawn conclusions:* Once the biphasic equation is made continuous, the two species differ by 1.13-fold at 240 h under version 1’s own parameters. The comparative conclusion of version 1 was produced by the discontinuity and is withdrawn. The reported persister fraction of 3.58% was an input, not a fit. The abstract of version 1 compared two parameters drawn from different equations and attached one species’ transition time to the other.

*Statistics:* The Kolmogorov-Smirnov test of version 1 is removed: applied to deterministic curves its p-value is set by the density of the simulation grid, falling by more than 170 orders of magnitude as the grid is refined. The reported R-squared above 0.9 is no longer used for model selection, because a structurally broken model and its correct replacement both exceed it on the same data while their corrected AIC differs by up to 84.

*Version 1’s experimental-data figure is withdrawn:* It was captioned “experimental data vs. model fitting”. No dataset was named in version 1 and none exists. The replacement figure in this version uses synthetic data with known ground truth, is labelled as such, and supports conclusions about the estimator only.

*Methods corrected:* Version 1 stated that differential equations were solved with LSODA and cross-validated with Euler integration. Every model in version 1 is a closed-form algebraic solution; nothing was integrated. That sentence is removed. The present version does contain a genuine initial value problem, the state-structured model of Section 2.3, and names the solver used for it.

*References:* All thirty references of version 1 were checked against PubMed. Seventeen verified, eleven required correction, and two could not be located and are removed. Version 1’s reference 3, the consensus definitions paper this work depends on, was cited with the wrong title, the wrong issue and an author who is not on it. The corrected list carries a PMID for every entry.

*What is added:* A state-structured replacement model in which biphasic killing emerges rather than being imposed, sterilisation time is finite, and the three survival strategies separate onto orthogonal measurable axes; a global sensitivity analysis; and an identifiability analysis of both the old and the new model. All code is public and regenerates every number in this version. Every number in this version is produced by code in the accompanying repository and can be regenerated with a single command. Numbers are cross-referenced to the output file that contains them.

## 1. Introduction

Antibiotic resistance, tolerance and persistence lead to treatment failure and prolonged infection [1,2]. Resistance is a heritable change that raises the concentration required to inhibit growth. Tolerance and persistence are non-inherited: a tolerant population dies more slowly at an unchanged minimum inhibitory concentration, and a persistent population contains a subpopulation that survives exposure that clears the bulk [3,4]. These are three different statements about three different measurements, and Brauner and colleagues set them out as such [4]: resistance is a change in the minimum inhibitory concentration (MIC), tolerance a change in the minimum duration for killing (MDK) at fixed MIC, and persistence a change in the shape of the killing curve rather than in either summary statistic.

*Staphylococcus aureus* and *Mycobacterium tuberculosis* sit at opposite ends of a growth-rate spectrum, with doubling times differing by roughly seventeen-fold, and they are treated on correspondingly different timescales [5,6]. *M. tuberculosis* persists in host tissue and requires months of therapy, a durability associated with dormancy and phenotypic heterogeneity [7,8,9,10]. *S. aureus* is a rapid grower whose biofilm and stationary-phase populations show marked tolerance and relapse after apparently successful treatment [11,12]. The contrast is a natural setting in which to ask what actually determines how long treatment must last.

Answering that question requires a model in which resistance, tolerance and persistence are different things. Much of the modelling literature, including our own earlier version of this work, does not meet that requirement. When all three are written as N(t) = N₀e^(−kt) with different values of k, they differ only in a number, and no measurement can distinguish them because the model contains no measurement that could. The problem is structural rather than numerical, and it cannot be fixed by re-estimating k.

This paper does three things. First, it establishes what goes wrong with the closed-form biphasic killing law that is commonly used to represent persistence, including a discontinuity that has apparently gone unremarked and a structural non-identifiability that makes its transition time uninterpretable. Second, it develops a state-structured replacement in which the three strategies occupy orthogonal directions in a space of measurable quantities. Third, it uses global sensitivity analysis of the replacement to identify which parameter actually governs time to sterilisation, and finds an answer different from the one a one-at-a-time analysis of the closed-form law returns.

We are explicit about the limits. This is a framework paper. It contains no experimental data, its parameter values are illustrative, and its species-level statements are therefore qualitative. What it does establish is which quantities must be measured, and what a model must contain before those measurements can be interpreted.

## 2. Methods

### 2.1 Notation and the models under examination

Let N(t) denote colony-forming units per millilitre at time t hours after the start of a constant antibiotic exposure, N₀ the inoculum, K the carrying capacity, r the replication rate, and C the drug concentration in multiples of a reference MIC.

**Logistic growth without drug.**

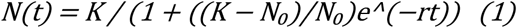

**The closed-form laws under examination.** We examined three closed-form expressions in common use, written here as they appear in the literature and in version 1 of this work:

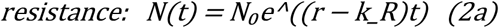

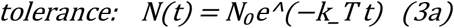

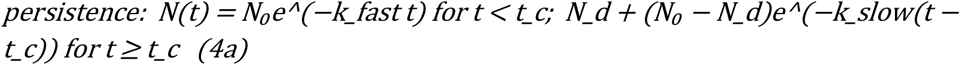

with N_d the dormant subpopulation and t_c the transition time. Section 3.1 to 3.3 establishes why each requires replacement.

**Corrected closed forms.** Equations 2a and 3a omit the carrying capacity declared in Equation 1 and, in the case of 3a, omit replication entirely. Both are replaced by the same differential form with the appropriate kill constant:

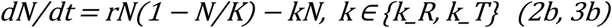

which has a closed solution: logistic with effective rate r − k and effective capacity K(r − k)/r.

Equation 4a is replaced by a biexponential, which is smooth everywhere, reaches sterilisation, keeps the dormant fraction f dimensionless, and has three rate parameters rather than four:

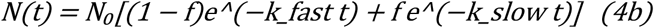

Under Equation 4b the transition time is an output rather than a parameter:

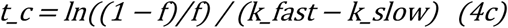

A continuous piecewise form, N(t ≥ t_c) = N₀e^(−k_fast t_c)e^(−k_slow(t − t_c)), is also examined in Section 3.1 as the minimum repair to Equation 4a. It removes the discontinuity but retains an imposed t_c and is therefore not recommended.

### 2.2 Pharmacodynamic function

None of Equations 2 to 4 contains a concentration term, so none can express an MIC, a dose or a regimen. We introduce a sigmoid maximum-effect function:

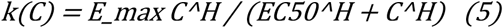

with E_max the maximum kill rate, EC50 the half-maximal concentration and H the Hill coefficient. This function is standard in antibacterial pharmacokinetic–pharmacodynamic modelling [13,14]. It is what makes the three survival strategies formally distinct, as set out in Section 2.4.

### 2.3 State-structured model

Replicating (S) and dormant (P) subpopulations exchange at first-order rates and each carries its own concentration–response:

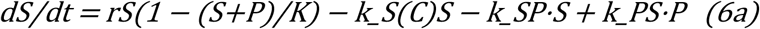

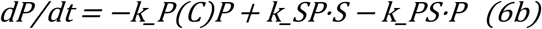

with k_S(C) and k_P(C) given by Equation 5 with compartment-specific E_max and EC50. In the absence of drug the dormant fraction equilibrates at k_SP/(k_SP + k_PS). Simulations start from that equilibrium, which is the consistent choice for an inoculum grown without drug.

Equations 6a and 6b contain no transition time. Biphasic killing arises because the two compartments are killed at different rates and the observable is their sum. The transition is recovered as a derived quantity, defined as the time of maximum curvature of log₁₀N(t) over the interval in which the population remains above the limit of detection.

Systems were integrated with the LSODA method (SciPy `solve_ivp`, relative tolerance 10⁻¹⁰, absolute tolerance 10⁻⁶). Unlike Equations 1 to 4, which are algebraic and require no solver, Equations 6a and 6b are a genuine initial value problem.

### 2.4 Encoding the three strategies

Within Equations 5 and 6 the three strategies are three different modifications of one parameter set:

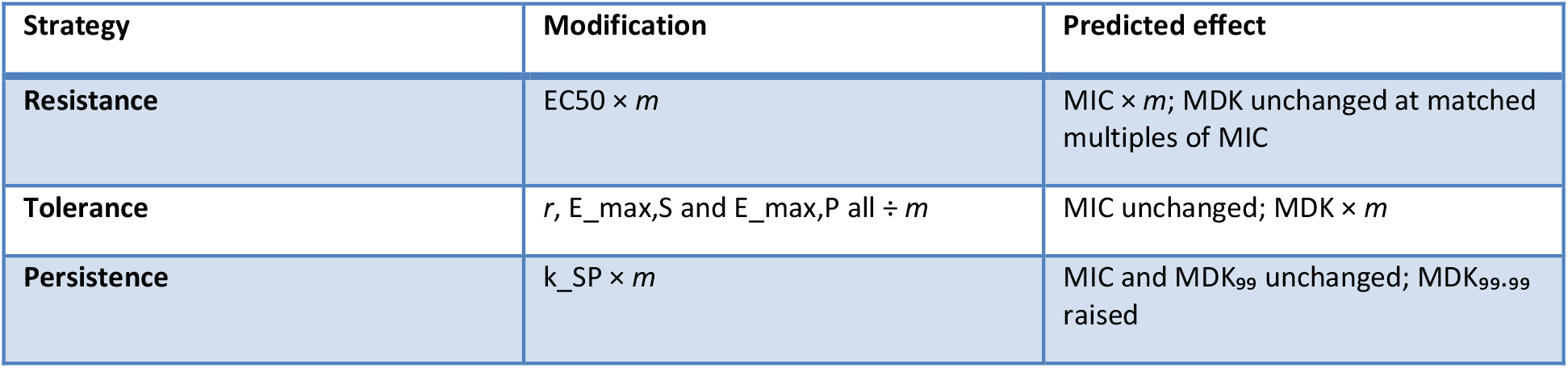

The encoding of tolerance requires comment. Dividing E_max alone is not tolerance. It lowers the maximum kill rate without slowing replication, which moves the balance point at which kill and growth cancel and therefore shifts the MIC by approximately *m*^(1/H). That is partial resistance. Tolerance is a slowing of metabolism, in which replication and killing scale together, and it is this joint scaling that leaves the MIC exactly invariant while multiplying the MDK. Conflating the two is precisely the confusion that a structured model exists to prevent.

### 2.5 Endpoints

All endpoints are computed from the same simulated curve.

- **MIC**: the lowest concentration at which the initial net growth rate of the bulk population is non-positive, located by Brent root-finding.
- **MDK₉₉ and MDK₉₉.₉₉**: the times at which the surviving fraction first reaches 10⁻² and 10⁻⁴ [4].
- **Time to sterilisation**: the time at which the population first falls below an assay limit of detection, taken as 10² CFU/mL, the value used in the hollow-fibre studies catalogued in the accompanying data manifest.
- **log₁₀ reduction at 240 h**.

Endpoints are located by bracketing followed by Brent root-finding on an interpolated curve. Where an endpoint is not reached, the search window is doubled repeatedly to 10⁶ h before infinity is returned, so a reported infinity means the endpoint is genuinely unreachable and never that the search was truncated. This distinction matters in Section 3.1.

### 2.6 Parameters

Parameter values used in this work fall into two classes, and the distinction is maintained throughout.

#### Class A, values from version 1’s parameter table

These are used only in Sections 3.1 to 3.4, where the object of study is the closed-form equations themselves. Every number in that table was adopted from the literature without a traceable extraction; none is a measurement tied to a named dataset, figure panel and digitisation record. They are reproduced here unchanged because the point of Sections 3.1 to 3.4 is that the equations misbehave *on their own stated parameters*, which requires using those parameters and no others.

| Parameter | *M. tuberculosis* | *S. aureus* |
| --- | --- | --- |
| Replication rate $r$ | 0.03 /h | 0.5 /h |
| Resistance kill rate $k_R$ | 0.002 /h | 0.01 /h |
| Tolerance kill rate $k_T$ | 0.05 /h | 0.2 /h |
| Fast kill rate $k_{fast}$ | 0.1 /h | 0.5 /h |
| Slow kill rate $k_{slow}$ | 0.001 /h | 0.01 /h |
| Dormant fraction $f$ | 1–5% | 0.01–0.1% |
| Asserted transition time $t_c$ | 80 h | 12 h |

#### Class B, illustrative values for the state-structured model

These were chosen to reproduce documented qualitative behaviour, namely a doubling time near 23 h with a deep non-replicating compartment for *M. tuberculosis* and near 1.4 h with a shallow, transient tolerant state for *S. aureus*. **They are not measurements.** They are labelled as illustrative on every figure in which they appear, and no quantitative species-level claim is made from them.

### 2.7 Estimation, identifiability and model comparison

Time-kill data are approximately homoscedastic on the log₁₀ scale, so all fitting was performed on log₁₀ CFU/mL by nonlinear least squares (SciPy `least_squares`, trust-region reflective). Observations below the limit of detection were treated as left-censored, contributing a residual only when the model predicted a value above the limit.

Because no experimental dataset exists for this work, fitting was performed on **synthetic** data generated from Equations 6a and 6b with known ground truth, additive Gaussian noise on log₁₀ CFU/mL, triplicate plating and censoring at the limit of detection. Two sampling designs were used per species, matched to that species’ kill window. This design supports two conclusions and no others: whether the estimator recovers the parameters that generated the data, and whether a given model’s parameters are identifiable from data of realistic density. It supports no conclusion about either organism, and every figure built on it is stamped accordingly.

Model quality was assessed by the corrected Akaike information criterion, the Bayesian information criterion, a Wald–Wolfowitz runs test on the residual sign sequence, and the condition number of the Jacobian at the optimum. Parameter identifiability was assessed by profile likelihood, refitting all other parameters at each fixed value and taking the 95% interval from the likelihood-ratio statistic against the χ²(1) quantile. A profile whose interval reaches the edge of the searched range indicates a parameter that the data does not determine.

We deliberately do not report the coefficient of determination as a model-selection statistic. Section 3.8 shows why.

### 2.8 Sensitivity analysis

Three analyses were run.

3. **One-at-a-time, ±10%, on the closed-form law**, reproducing the method used in version 1.
4. **One-at-a-time, ±10%, on the state-structured model**, so that method and model can be varied independently.
5. **Variance-based global analysis** of the state-structured model using the Saltelli estimator for first-order and total-order Sobol indices, with 512 base samples over seven parameters, giving 4,608 model evaluations per species. Parameters were drawn log-uniformly over ±50% of their illustrative values.

The gap between total-order and first-order indices measures the share of outcome variance arising from parameters acting jointly. A one-at-a-time analysis cannot produce this quantity at all, and where it is large, one-at-a-time conclusions are not merely imprecise but the wrong kind of statement.

### 2.9 Software and reproducibility

Python 3.14.6, NumPy 2.5.0, SciPy 1.18.0, pandas 3.0.3, Matplotlib 3.11.1. The full analysis runs in approximately 110 seconds on a laptop; the global sensitivity analysis accounts for about 92 s of that. All code, all intermediate tables and all figure scripts are in the accompanying repository. `python` `run_all.py` regenerates every number and every figure from scratch. Each stage writes a receipt recording library versions and run parameters.

## 3. Results

### 3.1 The closed-form biphasic law increases the population at its transition

Equation 4a is not a decay law. Evaluating its second branch at t = t_c gives

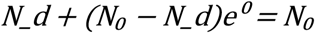

for any parameter values, so the modelled population returns to the inoculum at the transition irrespective of how much killing occurred in the first phase. The magnitude of the jump is the reciprocal of the first-phase survival.

| Parameter set | $N(t_c^-)/N_0$ | $N(t_c^+)/N_0$ | Upward jump |
| --- | --- | --- | --- |
| *M. tuberculosis*, $t_c = 80$ h | $3.36 \times 10^{-4}$ | 1.000 | $\times 2,981$ (3.47 $\log_{10}$) |
| *S. aureus*, $t_c = 12$ h | $2.48 \times 10^{-3}$ | 1.000 | $\times 403$ (2.61 $\log_{10}$) |
| *S. aureus*, $t_c = 80$ h | $4.25 \times 10^{-18}$ | 1.000 | $\times 2.35 \times 10^{17}$ (17.4 $\log_{10}$) |

*(’results/tables/eq4_discontinuity.csv’; Figure 1A-C)*

**Figure 1.**
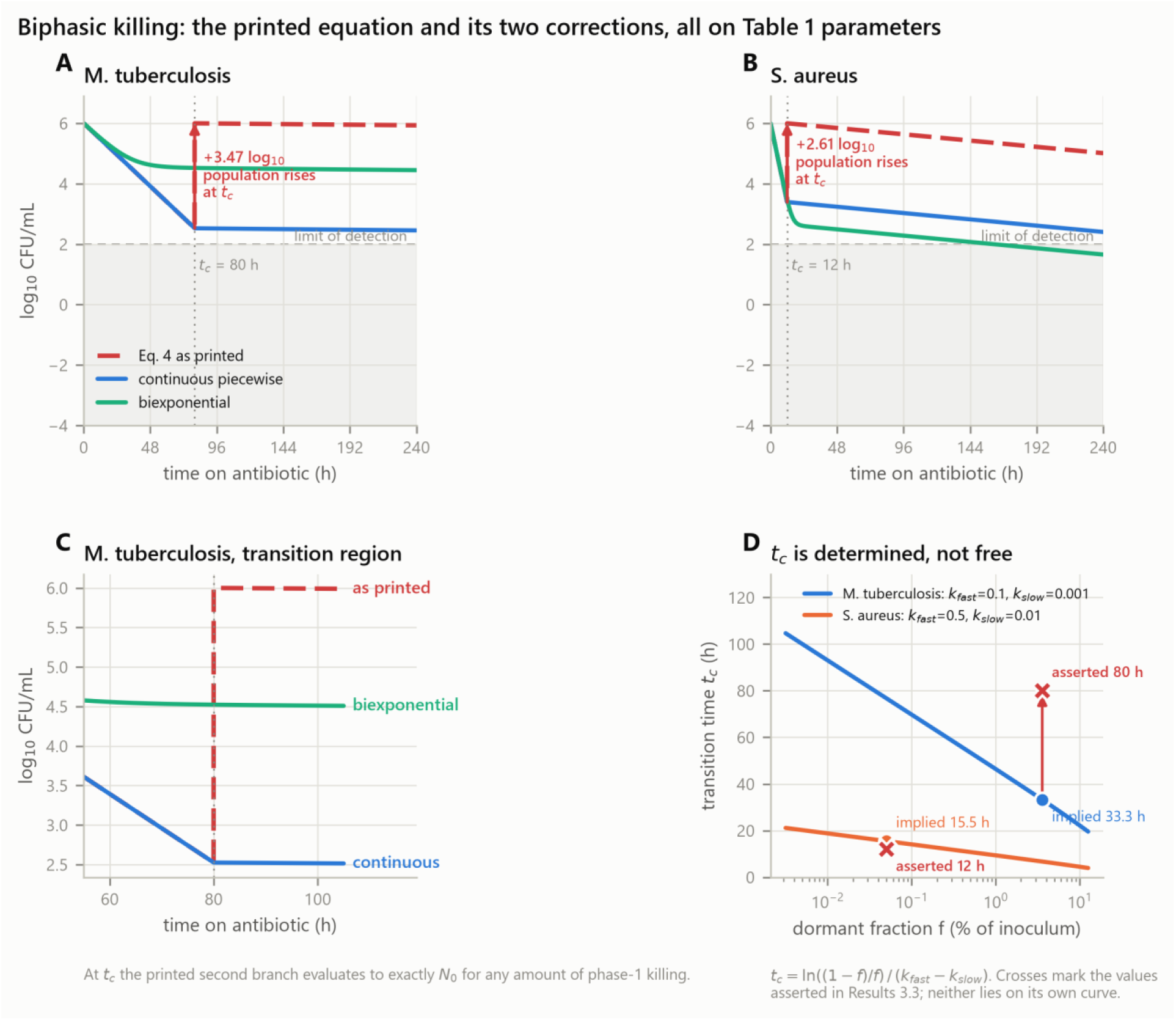
The closed-form biphasic law and its two corrections, all on the same parameters. (A, B) Equation 4a against the continuous piecewise and biexponential replacements. The arrow marks the upward jump at t_c: 3.47 log₁₀ for *M. tuberculosis*, 2.61 log₁₀ for *S. aureus*. (C) Detail of the transition region. At t_c the second branch of Equation 4a evaluates to exactly N₀ for any amount of first-phase killing. (D) The transition time is fixed by the other three parameters through Equation 4c. Crosses mark the values asserted in version 1; neither lies on its own curve.

A second consequence follows from the same expression. As t grows, N(t) → N_d. The dormant subpopulation never declines, so the model asserts that no regimen of any duration can sterilise. Under the parameters of Section 2.6 this places a permanent floor at 3.6 × 10⁴ CFU/mL for *M. tuberculosis*, which is 358 times the assay limit of detection. Time to sterilisation is therefore infinite, not as an artefact of a truncated search but as a property of the equation; the search window was extended to 10⁶ h before infinity was returned (`results/tables/model_endpoints.csv`).

Both corrected forms give finite sterilisation times under the same parameters: 1,290 h for *M. tuberculosis* and 333 h for *S. aureus* under the continuous piecewise form, and 5,881 h and 161 h under the biexponential.

A published figure drawn from Equation 4a would show a vertical rise of two to three log₁₀ at the breakpoint. Figures that do not show this were not generated by the equation their methods section states. This discrepancy must be checked and disclosed wherever the equation appears.

### 3.2 The transition time is determined by the other parameters

In a two-subpopulation system the crossover occurs where the two exponential terms are equal, at t_c = ln((1−f)/f)/(k_fast − k_slow) (Equation 4c). It is therefore not available as a free parameter.

| Species | $t_c$ asserted | $t_c$ implied by $k_{\text{fast}}$, $k_{\text{slow}}$, $f$ | Discrepancy |
| --- | --- | --- | --- |
| *M. tuberculosis* | 80 h | 33.3 h | 2.41× |
| *S. aureus* | 12 h | 15.5 h | 0.77× |

*(’results/tables/tc_identifiability.csv’; Figure 1D)*

Neither asserted value lies on its own curve. Treating f, k_fast, k_slow and t_c as four free parameters over-parameterises a three-parameter system, and Section 3.8 shows that the consequence is not theoretical: t_c cannot be estimated from time-kill data of realistic density.

### 3.3 The resistance and tolerance equations omit terms their own parameters require

Equation 2a has no carrying-capacity term although Equation 1 declares one. Over a 240 h window this produces 10⁵⁷ CFU/mL for *S. aureus*, which exceeds the estimated total prokaryotic population of Earth [15] by twenty-six orders of magnitude. Applying the carrying capacity already declared in Equation 1 (Equation 2b) removes the excursion (Figure 2).

**Figure 2.**
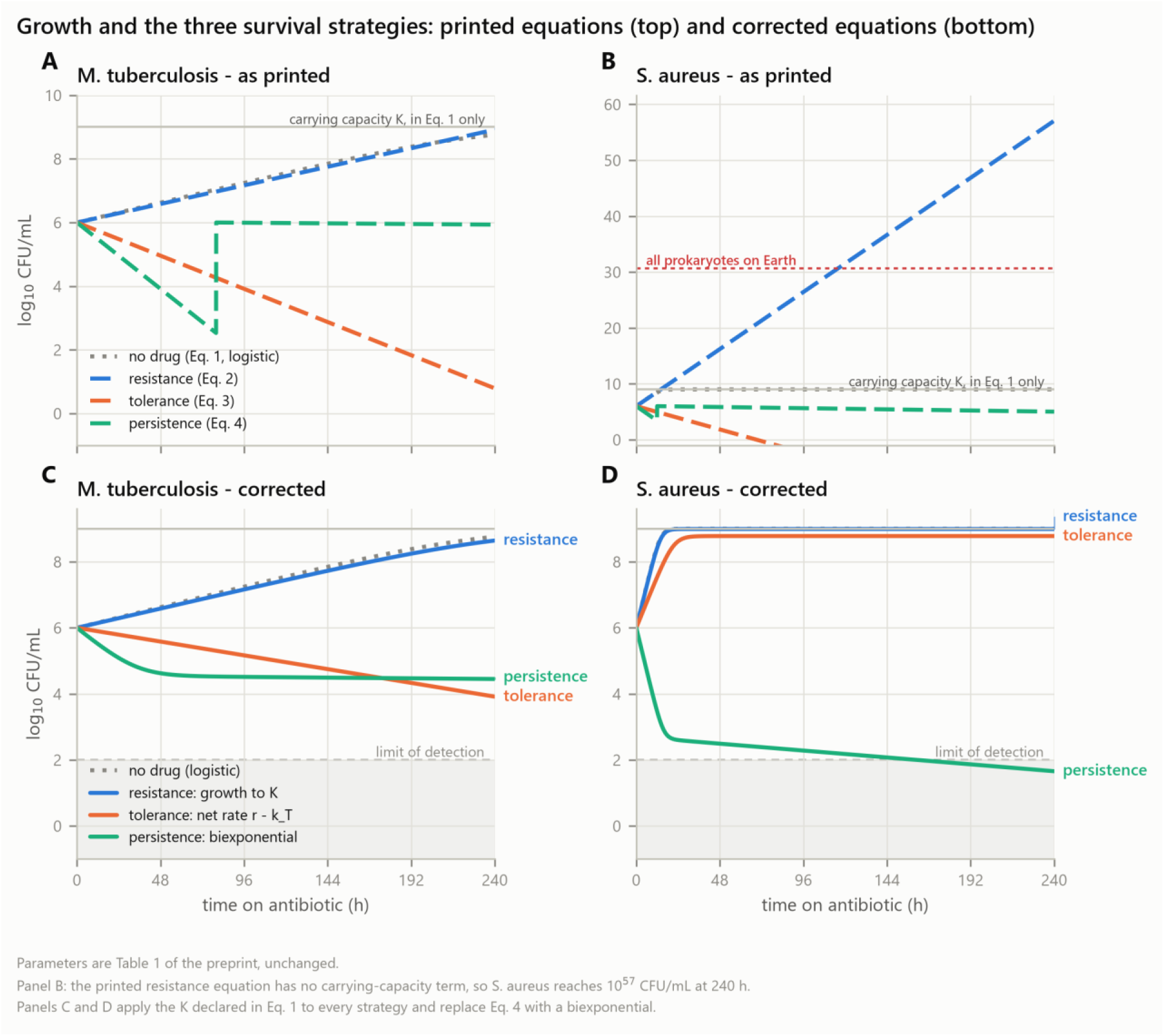
Growth and the three survival strategies, printed equations and corrected equations. (A, B) Equations 1, 2a, 3a and 4a on the Class A parameters of Section 2.6, unchanged. The carrying capacity declared in Equation 1 appears in none of the survival laws, so the resistance curve for *S. aureus* passes 10⁵⁷ CFU/mL by 240 h, exceeding the estimated prokaryotic population of Earth by twenty-six orders of magnitude. The vertical rise in the persistence curve is the discontinuity of Section 3.1. (C, D) The same scenarios with the carrying capacity applied to every strategy and Equation 4a replaced by the biexponential. Note that under the corrected tolerance equation the *S. aureus* population grows, because its replication rate exceeds its tolerance kill rate.

Equation 3a omits replication entirely, and for a fast grower this reverses the direction of the result rather than merely changing its magnitude. With r = 0.5/h and k_T = 0.2/h, the net rate is **+0.30 /h**: the population grows to carrying capacity. Equation 3a instead reports a decline of fifteen log₁₀ over the same window (`results/tables/eq3_omits_replication.csv`). For *M. tuberculosis* the sign is correct, because r = 0.03 < k_T = 0.05, but the magnitude is still wrong. A tolerance equation without a replication term cannot be applied to an organism whose replication rate appears in the same parameter table.

More fundamentally, Equations 2a and 3a are the same equation with different constants. As printed they do not distinguish resistance from tolerance, contradicting the framework they are meant to implement [4].

### 3.4 The comparative conclusion does not survive the correction

The strongest reason to abandon rather than patch Equation 4a is that the species contrast it produces is an artefact of its discontinuity. Correcting the equation to the continuous piecewise form and changing nothing else gives, at 240 h:

| Species | Surviving fraction, Eq. 4a | Surviving fraction, corrected |
| --- | --- | --- |
| *M. tuberculosis* | 0.857 | $2.86 \times 10^{-4}$ |
| *S. aureus* | 0.103 | $2.54 \times 10^{-4}$ |

The two species differ by a factor of **1.13, or 0.05 log₁₀**

(`results/tables/central_comparison.csv`; Figure 1A-B). On the parameters of Section 2.6 they are indistinguishable at ten days once the equation is made continuous. The apparent contrast in the corresponding figure of version 1 was produced by the fact that *M. tuberculosis*, with the later transition time, spent more of the window on the resurrected branch.

The biexponential form (Equation 4b) does preserve a difference, because it retains the dormant fraction as a weight on a surviving term rather than discarding it, and it is the form we recommend where a closed expression is required. But the general point stands: a conclusion that changes sign or magnitude when a discontinuity is removed was a property of the discontinuity. This is why the remainder of this paper proceeds mechanistically.

### 3.5 Biphasic killing emerges from compartment structure

Equations 6a and 6b contain no breakpoint, yet they produce biphasic kill curves. At four times the MIC and an inoculum of 10⁶ CFU/mL, the curvature maximum of the total population occurs at 25 h for the slow grower and 15 h for the fast grower, and the population reaches the limit of detection at 380 h and 21 h respectively (`results/tables/fig05_mechanistic.csv`, Figure 3).

**Figure 3.**
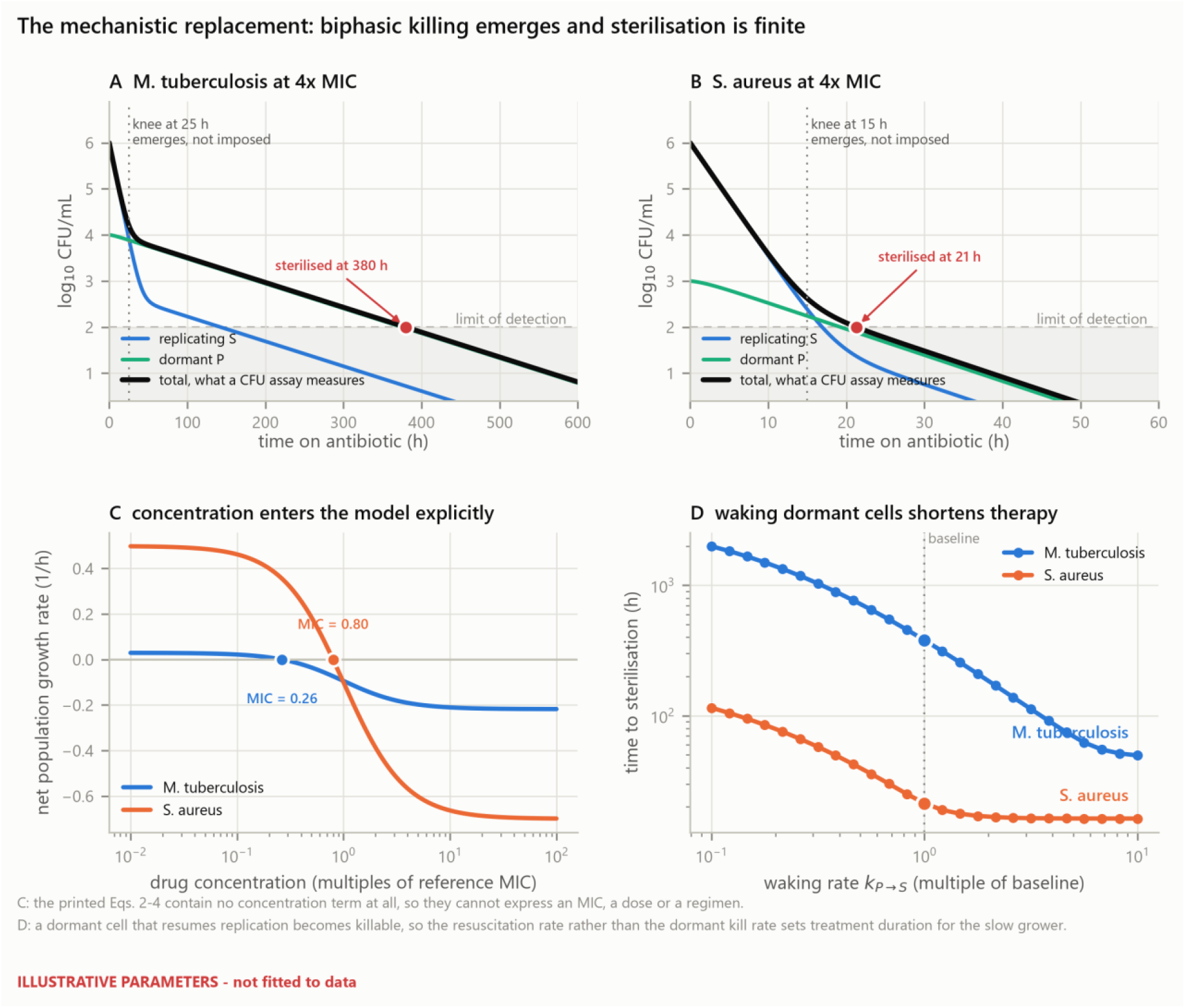
The state-structured model. (A, B) Replicating, dormant and total populations at four times MIC. Biphasic killing emerges from the compartment structure; the model contains no transition time, and the curvature maximum is a derived observable. Sterilisation occurs in finite time. (C) Net growth rate against concentration, with the MIC of each species marked. Equations 2a to 4a contain no concentration term and cannot produce this panel. (D) Time to sterilisation against the resuscitation rate. Illustrative parameters.

Three properties distinguish this from Equation 4a. The curve is continuous and differentiable throughout. Sterilisation occurs in finite time for any non-zero dormant kill rate, so treatment duration is a computed quantity rather than infinite by construction. And the transition is an observable derived from the rate constants rather than a parameter fitted independently of them, so it cannot contradict them in the way documented in Section 3.2.

The compartment trajectories also make the mechanism visible. The dormant compartment declines faster than its own kill rate alone would allow, because cells leaving dormancy enter a compartment that is being killed rapidly. The effective clearance rate of the dormant pool is approximately k_PS + E_max,P rather than E_max,P. This observation is developed in Section 3.7.

### 3.6 The three strategies occupy orthogonal directions

Introducing each mechanism separately into one parameter set, and measuring each strain at eight times its own MIC so that a resistant strain is not merely under-dosed:

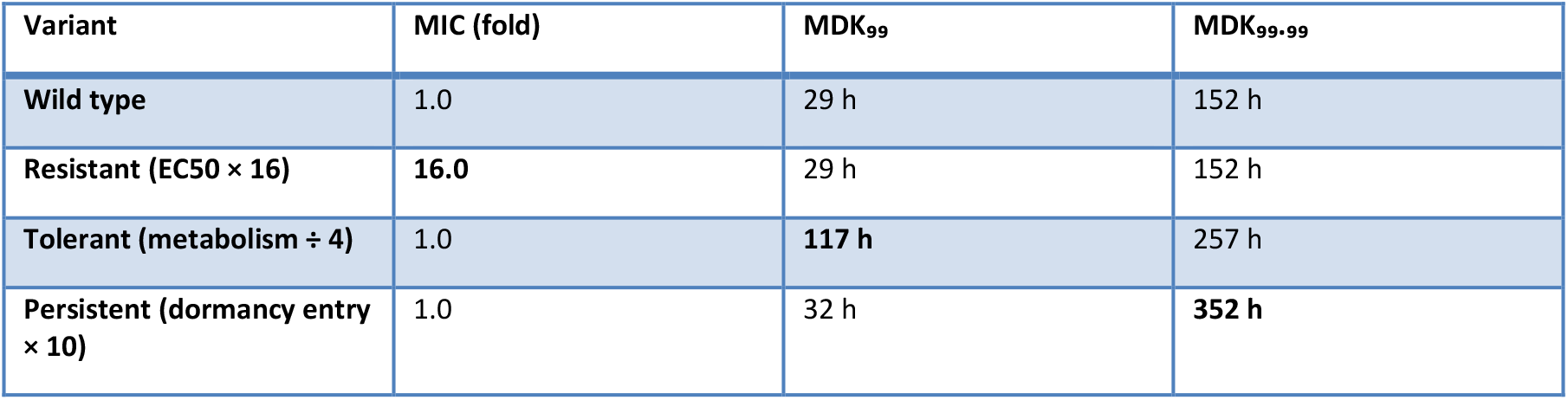

*(’results/tables/fig06_mic_mdk.csv’, Figure 4)*

**Figure 4.**
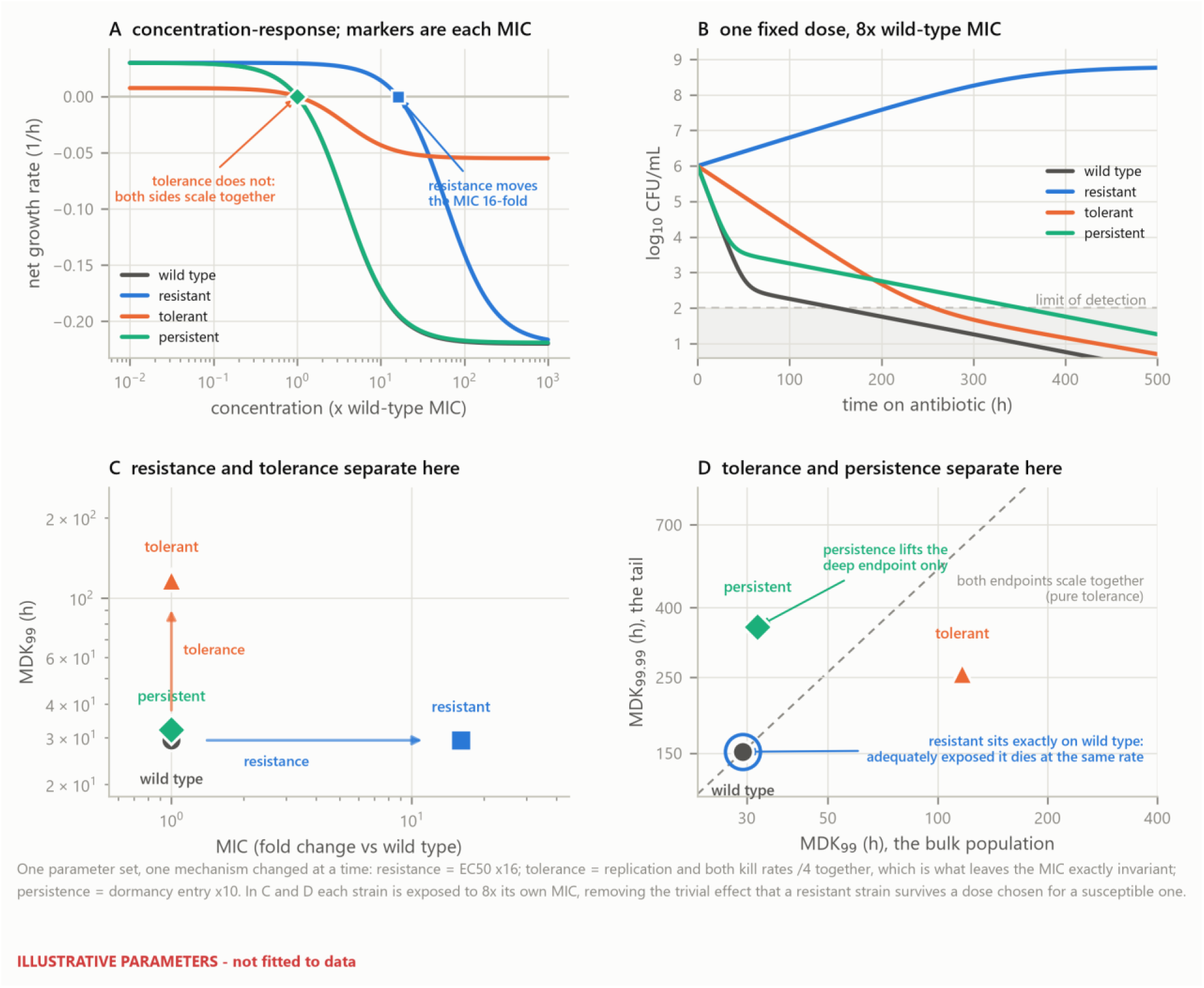
Three strategies, three signatures. One parameter set with one mechanism changed at a time. (A) Concentration–response; markers give each variant’s MIC. Resistance moves the MIC sixteen-fold; tolerance does not move it at all, because replication and killing scale together. (B) Time-kill at a single fixed dose of eight times the wild-type MIC, the clinically visible situation, in which the resistant variant grows. (C) The MIC–MDK₉₉ plane separates resistance from tolerance. (D) The MDK₉₉–MDK₉₉.₉₉ plane separates tolerance from persistence. In C and D each strain is exposed to eight times its own MIC. Illustrative parameters.

Resistance moves the MIC sixteen-fold and leaves both killing endpoints untouched: adequately exposed, a resistant strain dies at the wild-type rate. Tolerance leaves the MIC exactly unchanged and multiplies MDK₉₉ by 3.99, recovering the imposed four-fold slowing. Persistence leaves the MIC and MDK₉₉ essentially unchanged, moving MDK₉₉ by only 10%, while raising MDK₉₉.₉₉ 2.3-fold.

Two consequences are practical. An isolate can be placed in this space from two standard laboratory measurements, so the distinction is operational rather than verbal. And a study that reports only MIC and a single killing endpoint at a fixed dose cannot separate tolerance from persistence, because both raise the deep endpoint; the two are separated only by whether the bulk endpoint moves with it.

### 3.7 Resuscitation rate, not persister killing, governs sterilisation time

Applied to Equation 4a, one-at-a-time sensitivity analysis returns a ranking that is an algebraic identity rather than a result. For any t > t_c the second branch of Equation 4a contains k_slow and f and nothing else, so k_fast and k_T have elasticities of exactly zero on any endpoint evaluated after the transition, and k_slow necessarily ranks first (`results/tables/analytic_elasticities.csv`; Figure 5A). This ranking would be unchanged for any parameter values and any organism, and therefore cannot support a biological conclusion.

**Figure 5.**
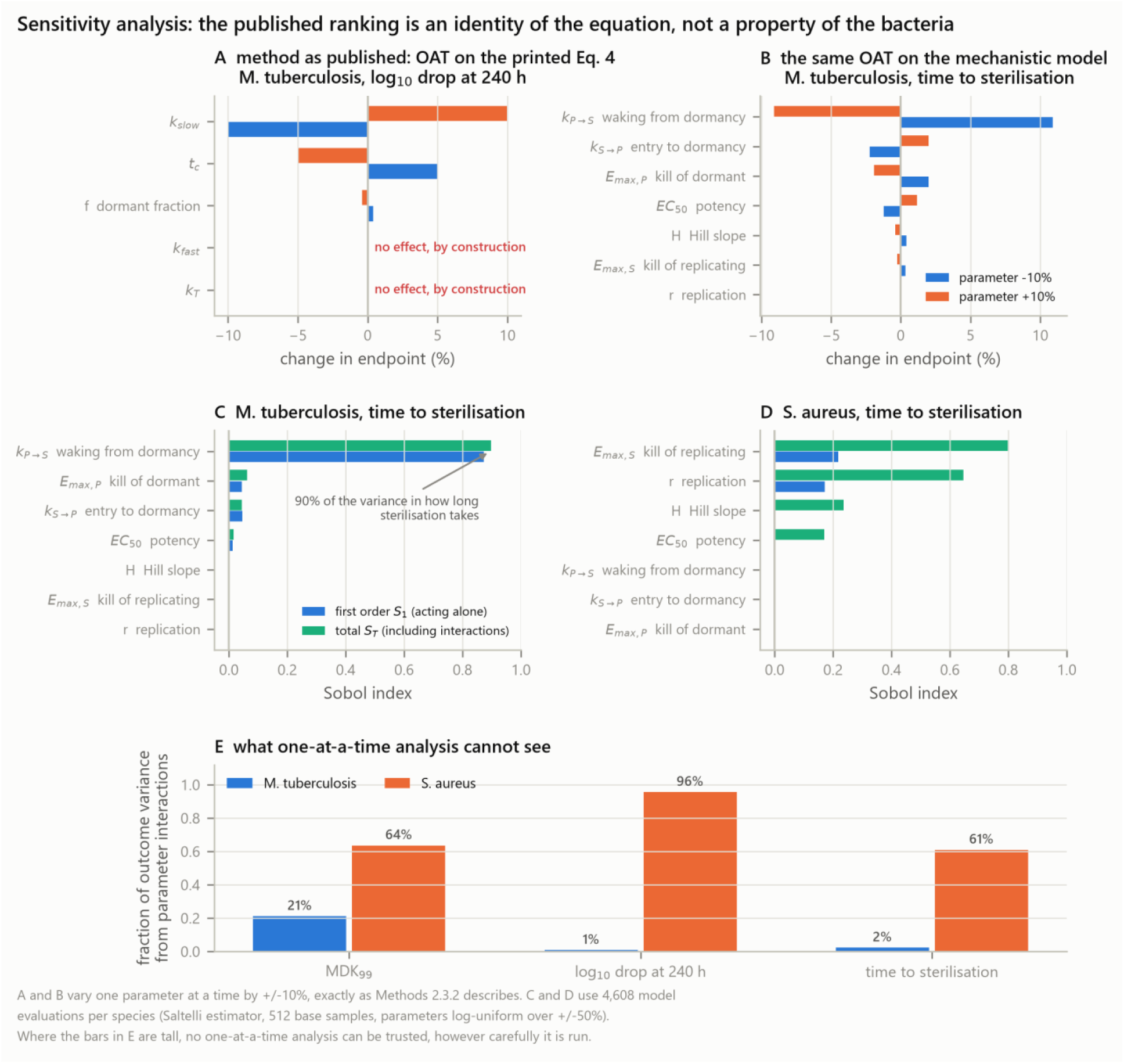
Sensitivity analysis: method and model varied independently. (A) One-at-a-time analysis at ±10% applied to Equation 4a. The elasticities of k_fast and k_T are exactly zero because neither appears in the branch that governs the endpoint. (B) The same analysis on the state-structured model, where every parameter acts at every time. (C, D) First-order and total-order Sobol indices for time to sterilisation, 4,608 model evaluations per species. (E) Share of outcome variance arising from parameter interactions. Where these bars are tall, one-at-a-time analysis is the wrong instrument.

Run on the state-structured model, where every parameter acts at every time, the answer is different. For the slow grower, the first-order Sobol index for k_PS, the rate at which dormant cells resume replication, is **0.873**, with a total-order index of **0.897** (`results/tables/sensitivity_sobol.csv`, Figure 5C). No other parameter reaches a total-order index of 0.07. Varying k_PS over the ±50% range shifts time to sterilisation from 380 h at baseline to 1,995 h at one tenth of baseline and 50 h at ten times baseline, a span of forty-fold.

The mechanism is the one identified in Section 3.5. A dormant cell is hard to kill; a dormant cell that resumes replication is not. Anything that increases the rate of resuscitation moves cells from a refractory compartment into a susceptible one, and the drug does the rest. This places resuscitation, rather than direct killing of persisters, at the centre of regimen shortening for slow-growing organisms, and it is a statement about a drug target that Equation 4a is structurally incapable of making.

For the fast grower the picture differs in kind, not just in detail. The dominant parameters are E_max,S and r, and **61% of the variance in time to sterilisation arises from parameter interactions** rather than from parameters acting alone; for the log₁₀ reduction at 240 h the interaction share reaches 96% (`results/tables/sensitivity_interaction_fraction.csv`, Figure 5E). Where the interaction share is that high, no one-at-a-time analysis of that organism can be trusted regardless of how carefully it is conducted.

### 3.8 The coefficient of determination cannot distinguish these models

Three further statistical claims from version 1 fail the same way, and are examined in Figure S1. Fitting Equation 4a and Equation 4b to the same synthetic time-kill data at four sampling designs produces R² above 0.9 for both models in three of the four designs, with a maximum difference between the two models of 0.098. Over the same fits the corrected Akaike information criterion differs by up to **84 units**, always favouring the biexponential (`results/tables/r2_vs_aicc_discrimination.csv`, Figure 6C).

**Figure 6.**
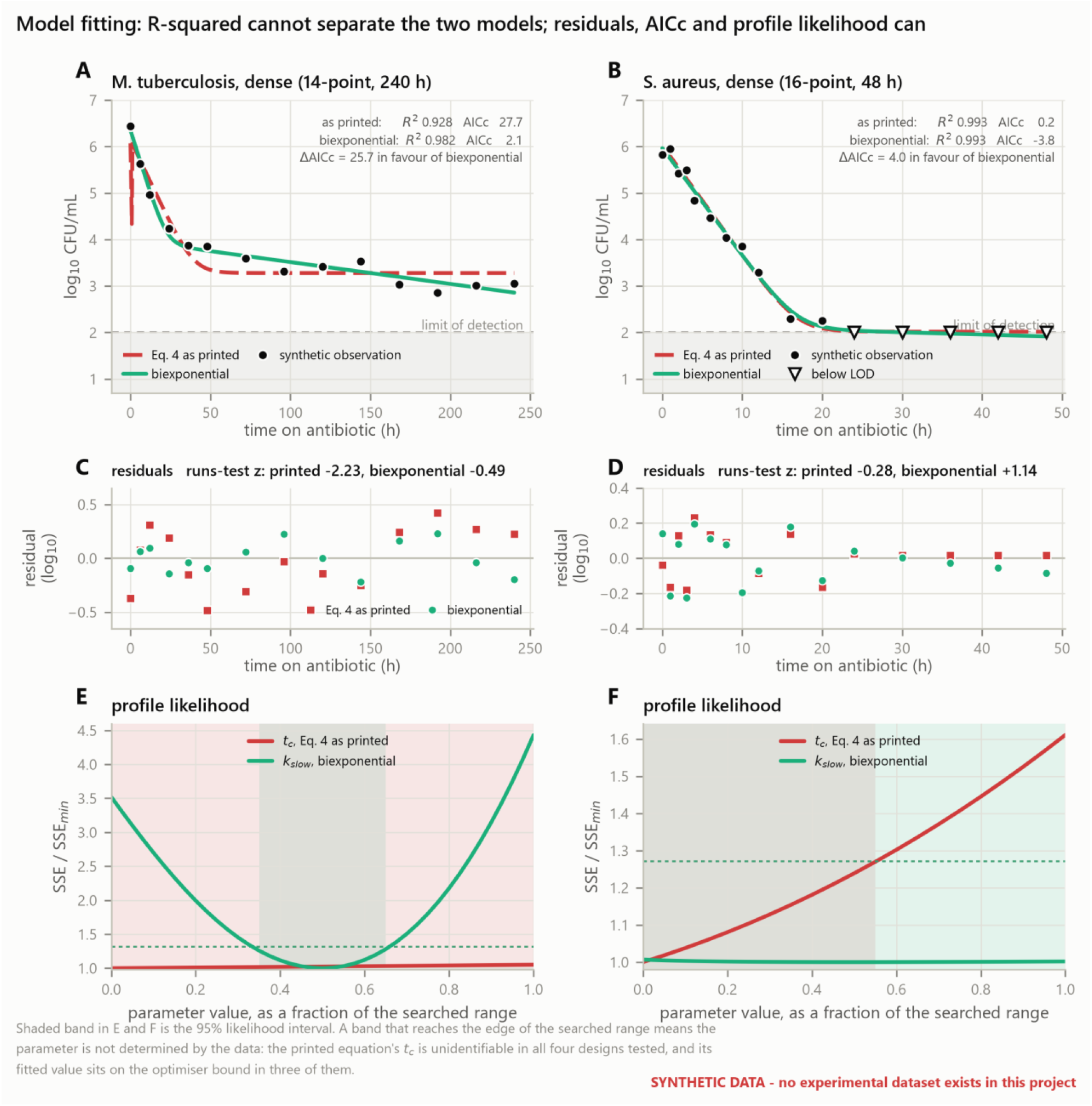
Fitting, model comparison and identifiability. Synthetic data with known ground truth; no experimental dataset exists for this work. (A, B) Both models track the data closely. (C, D) Residuals and runs-test statistics. (E, F) Profile likelihood. The 95% interval for the transition time of Equation 4a reaches the edge of the searched range in every design; the slow rate of the biexponential is identifiable for the slow grower.

**Figure S1.**
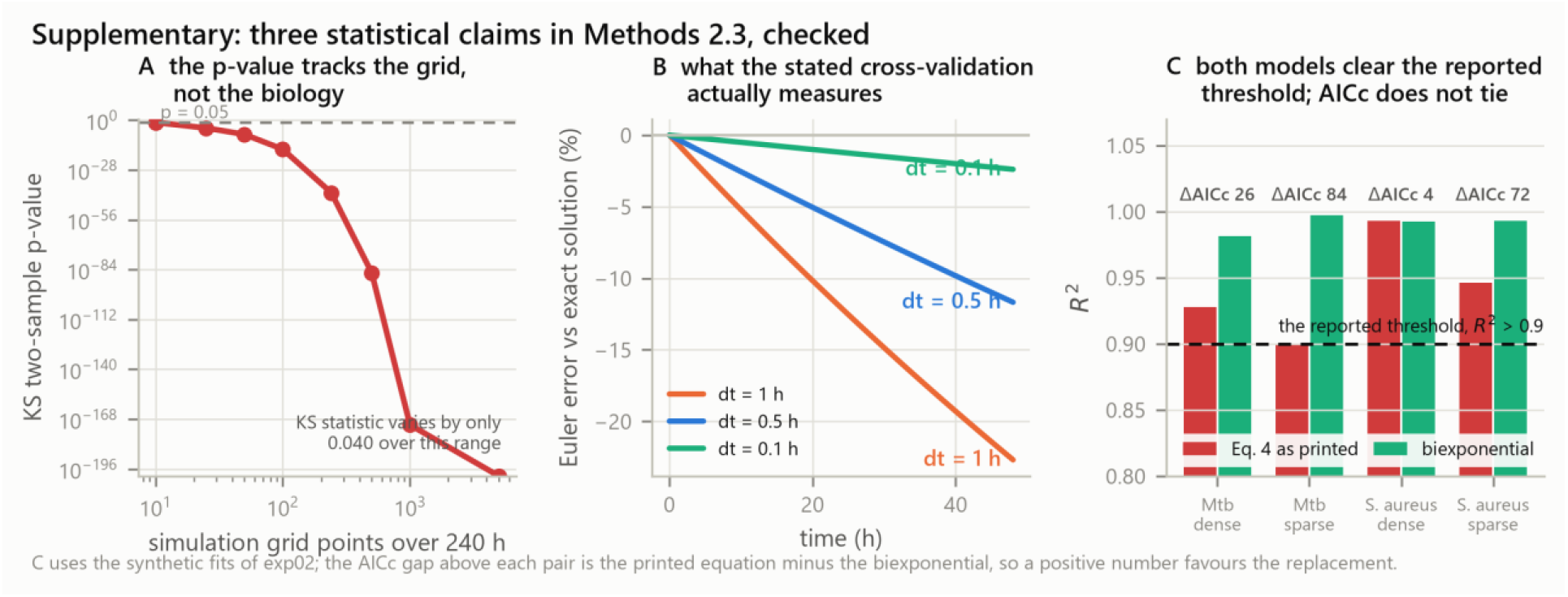
Three statistical claims checked. (A) The two-sample Kolmogorov–Smirnov p-value computed between two deterministic curves falls by more than 170 orders of magnitude as the simulation grid is refined, while the test statistic moves by 0.04. The test has no inferential content when applied to deterministic curves. (B) Euler integration error against the exact solution, which is what a comparison between Euler and a closed-form expression measures. (C) R² for both models against the frequently reported 0.9 threshold, annotated with the corrected Akaike information criterion difference.

R² is close to one for almost any decreasing function fitted to a monotone decaying curve, because the total sum of squares is dominated by the spread of the data across orders of magnitude. It is therefore compatible with a model that raises the population by three log₁₀ at its transition. It should not be used to select between candidate killing models, and a reported R² above 0.9 provides no evidence that a persistence model is correct.

Profile likelihood on the same fits shows that the transition time of Equation 4a is not identifiable. Its 95% likelihood interval reaches the edge of the searched range in **four of four designs**, including the two dense designs, and the point estimate lies on the optimiser bound in three of them (`results/tables/profile_tc.csv`, Figure 6E and 6F). The Jacobian condition number at the optimum reaches 5 × 10¹⁷ for Equation 4a against approximately 5 × 10² for Equation 4b, a difference of fifteen orders of magnitude that indicates a rank-deficient design matrix rather than a difficult optimisation.

A reported value of t_c from a fit of Equation 4a therefore carries no information about the organism. It records where the optimiser stopped.

## 4. Discussion

### 4.1 What this work establishes

Three results are independent of any parameter choice, because they are properties of the equations. The closed-form biphasic killing law increases the population at its transition and converges to an immortal floor. Its transition time is determined by its other three parameters and, in practice, is not estimable from time-kill data of realistic density. And a model in which resistance, tolerance and persistence are one exponential with three different rate constants cannot distinguish them, because it contains no quantity that differs between them.

One result depends on the model structure but not on the specific parameter values: given a two-compartment model with a concentration–response, the three strategies separate onto orthogonal axes of measurable quantities, and an isolate can be placed in that space from measurements laboratories already make.

One result depends on the illustrative parameters and is therefore provisional: the dominance of the resuscitation rate in setting time to sterilisation for the slow grower. Its mechanism is robust, in that any model with a refractory compartment and a susceptible compartment will show that moving cells between them changes clearance time. Its magnitude is not established and awaits fitting to data.

### 4.2 Implications, stated at the strength the evidence supports

If the dominance of resuscitation survives fitting to real time-kill data, it has a direct therapeutic reading: for slow-growing organisms, agents that drive dormant cells back into replication would shorten therapy more effectively than agents that kill dormant cells directly. This is consistent with the interest in resuscitation-promoting approaches and with the observation that treatment duration for tuberculosis is set by a small, slowly cleared subpopulation rather than by the bulk [7,8,9]. We emphasise that our analysis motivates this hypothesis rather than confirming it.

The measurement implication is firmer and does not depend on the parameter values. A study reporting only MIC and a single killing endpoint cannot distinguish tolerance from persistence. Separating them requires two killing endpoints at different depths, MDK₉₉ and MDK₉₉.₉₉, measured at matched multiples of each strain’s own MIC. Studies that dose all strains at a fixed absolute concentration measure under-dosing of resistant strains rather than any difference in their killing kinetics.

The methodological implication is firmest of all. A high R² is not evidence that a killing model is correct, and a fitted transition time from an over-parameterised biphasic law is not a measurement. Papers reporting either should also report an information criterion, a residual diagnostic and a profile likelihood.

### 4.3 Limitations

#### No experimental data

This is the principal limitation and it bounds everything else. Version 1 of this work presented a figure captioned as experimental data versus model fitting without identifying any dataset. That figure has been withdrawn. The present Figure 6 uses synthetic data with known ground truth, is stamped as such, and supports conclusions about the estimator and about identifiability only.

#### Illustrative parameters

The Class B values of Section 2.6 reproduce documented qualitative behaviour but are not measurements. No quantitative species-level claim is made from them. Candidate datasets suitable for fitting, principally hollow-fibre infection model studies for both organisms, are catalogued in the accompanying data manifest, and fitting the model to them requires no change to the model code.

#### Constant exposure

All simulations use a constant concentration. Real regimens produce fluctuating concentrations, and the relevant exposure metric may be the area under the curve relative to MIC, the peak relative to MIC or the time above MIC depending on drug class. Extending Equations 6a and 6b to time-varying C is straightforward and is the natural next step.

#### Two compartments

Dormancy is a continuum rather than a binary state, and depth of dormancy varies [16]. A two-compartment model is the simplest structure that separates the three strategies; it is not proposed as a complete description.

#### Deterministic

At the low copy numbers that determine sterilisation, extinction is a stochastic event and a deterministic model cannot give an extinction probability. A stochastic implementation is required for that question.

#### Sensitivity ranges

The global analysis samples ±50% log-uniformly around illustrative values. Once parameters are estimated, it should be rerun over the posterior.

### 4.4 Next steps

The binding constraint is data extraction rather than model development. In order: extract time-kill data from the identified hollow-fibre studies with a documented and versioned digitisation workflow; fit Equations 5 and 6 with a hold-out set designated before any fitting begins; recompute the global sensitivity analysis over the fitted posterior; and only then restate the species-level conclusions quantitatively. Extension to time-varying exposure and to a stochastic implementation follows.

## 5. Conclusion

Resistance, tolerance and persistence are three different phenomena, and a model that writes all three as one exponential with a different constant cannot tell them apart. The closed-form biphasic killing law widely used to represent persistence fails in three further ways that we quantify here: it raises the modelled population by two to three log₁₀ at its transition, it converges to a floor at which sterilisation is impossible at any duration, and its transition time is not identifiable from time-kill data of realistic density.

A two-compartment model with a sigmoid concentration–response removes all three failures and separates the strategies onto orthogonal axes of quantities that laboratories already measure. Applied to that model, global sensitivity analysis identifies the rate at which dormant cells resume replication as the dominant determinant of sterilisation time for a slow-growing organism, a determinant the closed-form law cannot express.

This is a framework rather than a measurement. Its quantitative claims about *M. tuberculosis* and *S. aureus* await fitting to experimental time-kill data. What it establishes is which quantities must be measured, and what a model must contain before those measurements mean anything.

## Data and code availability

All code, intermediate tables, receipts and figure-generating scripts are openly available at https://github.com/piranfar/comparative_model_presistance (code under the MIT licence, documentation and figures under CC BY 4.0). `python` `run_all.py` regenerates every number and every figure reported here in approximately 110 seconds, with no network access. Each analysis stage writes a receipt recording library versions and run parameters. No experimental dataset is used or distributed; candidate datasets for the fitting stage are catalogued with digital object identifiers in `data/manifests/datasets.csv`.

## Author contributions

V.P. conceived the study, wrote the code, performed the analyses and wrote the manuscript.

## Acknowledgements

To be completed by the author before submission.

**Candidate datasets named in Section 4.3 for the fitting stage.** These are cited as sources of data, not as support for any claim made here.

