## Supplementary Figures for "A state-structured pharmacodynamic framework separating resistance, tolerance and persistence, applied to Mycobacterium tuberculosis and Staphylococcus aureus"

### Supplementary: three statistical claims in Methods 2.3, checked

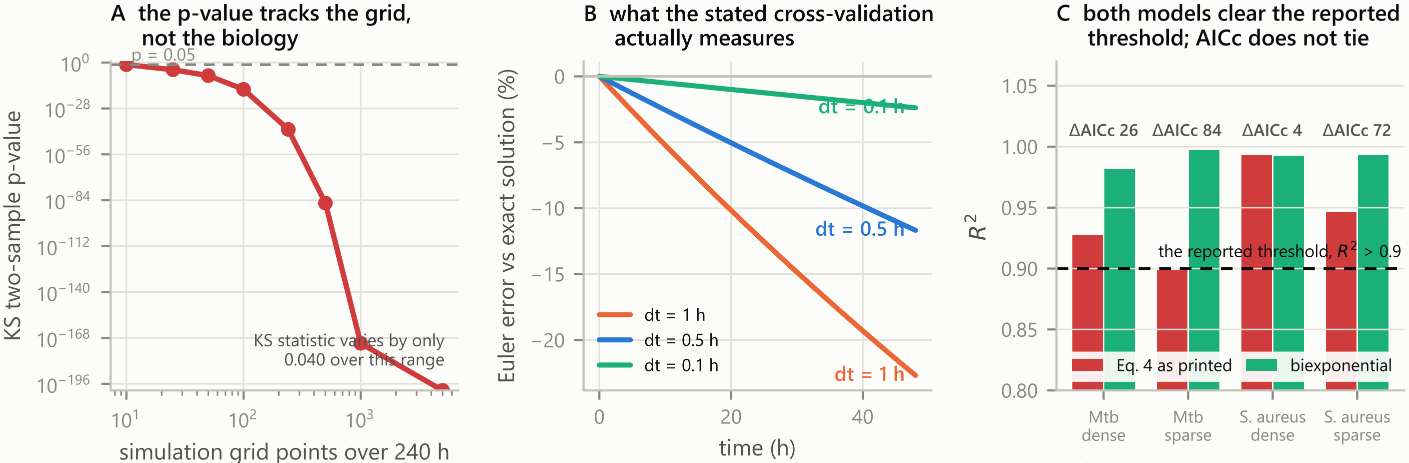


C uses the synthetic fits of exp02; the AICc gap above each pair is the printed equation minus the biexponential, so a positive number favours the replacement.
